# Three-dimensional structure of inner ear hair cell mitochondrial networks and ribbon synapses in a zebrafish model of Usher syndrome type 1B

**DOI:** 10.1101/2022.10.01.510403

**Authors:** Kenneth C. Riley, Alaa Koleilat, Joseph Dugdale, Shawna A. Cooper, Trace A. Christensen, Lisa A. Schimmenti

**Affiliations:** Department of Clinical Genomics, Mayo Clinic; Department of Laboratory Medicine and Pathology, Mayo Clinic; Department of Otorhinolaryngology, Head and Neck Surgery, Mayo Clinic; Mayo Clinic Graduate School of Biomedical Sciences, Mayo Clinic; Department of Biochemistry and Molecular Biology, Mayo Clinic; Microscopy and Cell Analysis Core, Mayo Clinic; Department of Ophthalmology, Mayo Clinic

**Keywords:** serial block-face scanning electron microscopy, zebrafish, hair cells, ribbon synapse, mitochondria

## Abstract

Inner ear hair cells are the fundamental unit of sound and vibration detection in inner ear and lateral line structures. Our understanding of the structure and ultrastructure of hair cells has heretofore relied upon two-dimensional imaging. The development of serial block-face scanning electron microscopy (SBFSEM) changes this paradigm and allows for ultrastructural evaluation of hair cells in three-dimensions. This is the first report of SBFSEM analysis in the *myo7aa*^*-/-*^ mutant where we evaluated several attributes of zebrafish hair cells from the inner ear cristae in both wildtype and *myo7aa*^*-/-*^ mutant zebrafish through three-dimensional reconstruction. We describe ribbon synapse number, location, and volume, mitochondrial localization, and innervation for individual hair cells. We determined that *myo7aa*^*-/-*^ mutant ribbon synapses have a smaller volume and surface area; however, all other hair cell attributes investigated were not significantly different between the mutant and wildtype zebrafish. These findings are critical for the development of therapies for deafness caused by mutations in *myo7aa*, as this study supports that the necessary hair cell machinery for hearing is largely intact. In addition, the methodology and measurements developed in this study provide a guide for the evaluation of zebrafish hair cells using SBFSEM.

## Introduction

Hair cells are specialized sensory cells that transduce vibration in air and water. In *Danio rerio* and other otophysian fishes, hair cells are located in multiple anatomic locations. For example, the inner ear, which consists of three apical cristae and otolith associated structures, specifically the utricle, saccule, and lagena^1^ **(Fig. 1)**. Additionally, the zebrafish lateral line, which contains mechanosensory hair cells arranged into clusters called neuromasts to detect water movement and vibration, depends on hair cells as well^2^. Starting at 5 days post-fertilization (dpf), zebrafish embryos are able to respond to sound in a reproducible and predictable manner^3^ with sound detection occurring in the saccule of the inner ear^4^.

**FIG. 1:**
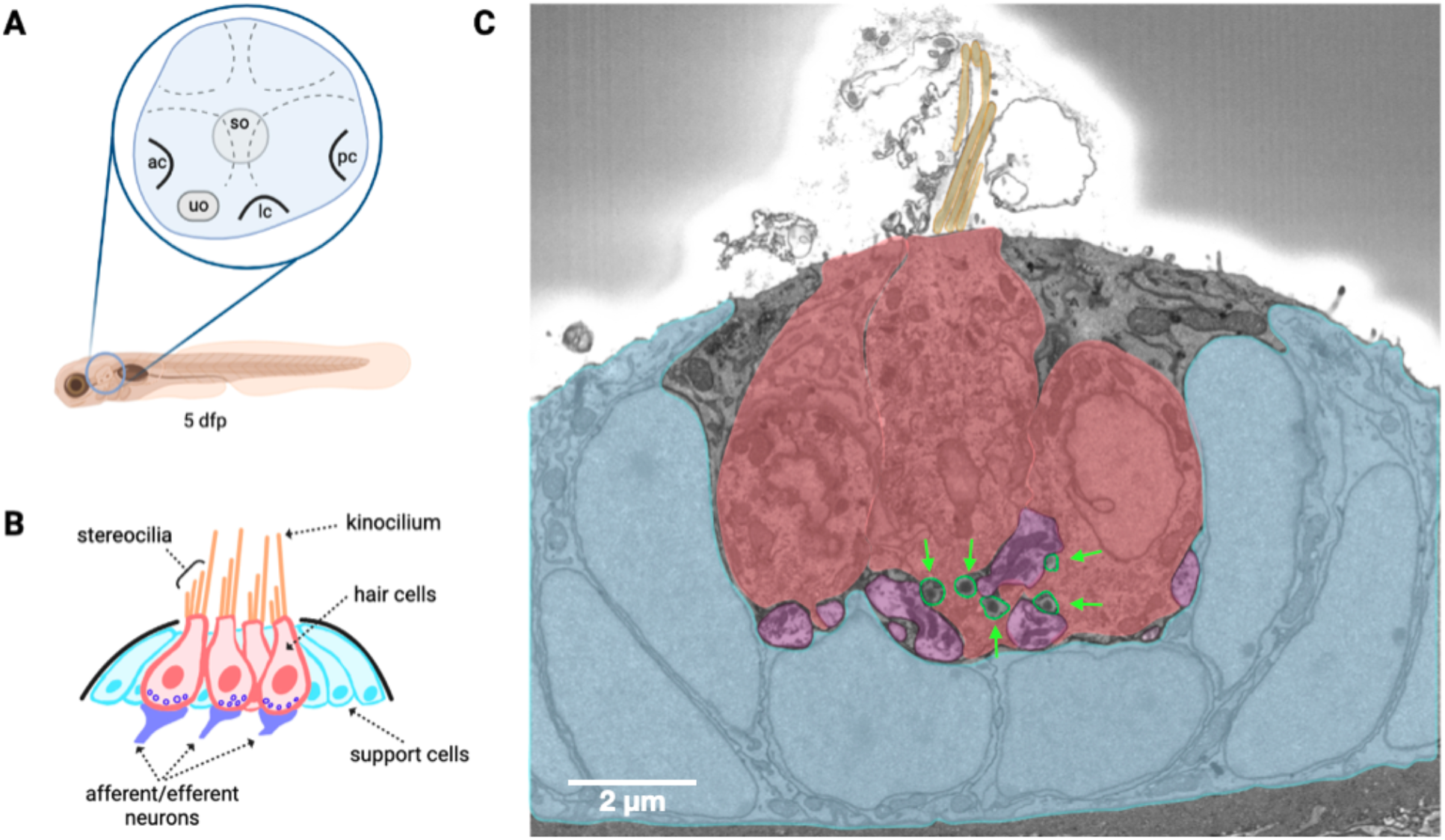
**(A)** Graphical depiction of 5 dpf zebrafish inner ear anatomy (lateral view). Dotted lines indicate epithelial columns, which form developing semicircular canals. Three apical cristae are shown: the anterior crista (ac), the lateral crista (lc), and the posterior crista (pc). The utricular otolith (uo) and saccular otolith (so) both lie atop macula, which are not shown. **(B)** Drawing of an apical crista. The cilia protruding from the superior region of the hair cells, stereocilia and kinocilia, deflect in response to water vibrations. This deflection leads to the transduction of vibrations into a sensory signal communicated via afferent neurons. This process is mediated by ribbon synapses, shown in small purple circles near the synapse. Created with BioRender.com. **(C)** Single SBFSEM image of a wildtype inner ear crista. Distinct features are masked in colors corresponding to the apical crista drawing: stereocilia (yellow), hair cells (red), the support cell (blue), and the afferent/efferent neurons (purple). Ribbon synapses are indicated by green arrows.

The complex structure of hair cells has largely been appreciated using various microscopy techniques that interrogate two-dimensional structure or ultrastructure^5^. However, there are limited reports of hair cell structures in the X, Y, and Z planes, which allow for a greater understanding of the position of organelles within the hair cell, their relationship to each other, and to the ribbon synapses^6-8^. The development of serial block-face scanning electron microscopy (SBFSEM) has now made it feasible to perform ultrastructural imaging of hair cells in all three dimensions allowing the investigator to visualize the interactions of organelles in ways that were not possible with two-dimensional imaging techniques^9^.

The *myo7aa*^*-/-*^ mutant zebrafish exhibits deafness and aberrant movement due to improper gating of the mechanotransduction channel responsible for depolarizing sensory hair cells and shows severe splaying of stereocilia within sensory hair cell bundles^10,11^. Furthermore, the *myo7aa*^*-/-*^ mutant is regarded as an ideal model of Usher syndrome type 1B in humans (USH1B)^11^, which is the most severe form of Usher syndrome characterized by irreversible, congenital, bilateral deafness and late-onset vision impairment^12^. The three-dimensional cellular ultrastructure in sensory hair cells of pathogenic variants in *myo7aa* is currently uncharacterized; however, this information may give clinically relevant insight to possible therapeutic targets.

It has been shown that *myo7aa*^*-/-*^ mutant neuromasts have fewer postsynaptic densities and lower numbers of ribbon synapses per lateral line sensory hair cell^13^. Moreover, mechanotransduction loss has been shown to not influence neuromast ribbon synapse localization or size in *sputnik* mutants, which carry a mutation in cadherin 23^14^. Both cadherin 23 and myosin VIIA act as structural proteins which help control the resting tension of tip-links between adjacent stereocilia; moreover, both *myo7aa*^*-/-*^ mutants and *sputnik* mutants lack mechanotransduction^11,14^. Therefore, we hypothesize the *myo7aa*^*-/-*^ mutants will present similar results within the inner ear cristae showing a decrease in the number of ribbons per hair cell and no change in ribbon localization, and stabilization.

In this report, we compared the three-dimensional ultrastructure of inner ear cristae hair cells in wildtype to *myo7aa*^*-/-*^ 5 dpf zebrafish. This is the first study to use SBFSEM to describe ribbon synapse number, volume and location in wildtype hair cells compared to *myo7aa*^*-/-*^ mutant hair cells. Additionally, we analyzed tethered synaptic vesicle organization, and qualitatively described mitochondrial network organization and hair cell innervation.

## Methods

### Zebrafish maintenance and husbandry

Animals used in this study were *Danio rerio* and strains used for experiments were *mariner* tc320 allele (c.2699T>A;p.Tyr846Stop) in exon 21 of *myo7aa*^11^. Adult fish were used to generate larvae, and control larvae were wildtype siblings of the *mariner* line. Animals were raised as per Koleilat *et al* ^13^. The Mayo Clinic Institutional Animal Care and Use Committee (IACUC) approved all experimental procedures.

### Serial block-face scanning electron microscopy and sample preparation

Samples for SBFSEM were prepared as previously described^15^. Briefly, fresh samples were fixed by immersion in 2% glutaraldehyde + 2% paraformaldehyde in 0.1 M cacodylate buffer containing 2 mM calcium chloride. After fixation, samples were rinsed in 0.1 M cacodylate buffer and placed into 2% osmium tetroxide + 1.5% potassium ferricyanide in 0.1 M cacodylate, washed with deionized and distilled (dd) water, incubated at 50°C in 1% thiocarbohydrazide, rinsed in dd water and placed in 2% uranyl acetate overnight. Next, the sample is rinsed in dd water, incubated with Walton’s lead aspartate, dehydrated through an ethanol series, and embedded in Embed 812 resin. To prepare the embedded sample for placement into the SBFSEM, a ∼1.0 mm^3^ piece was trimmed of any excess resin and mounted to an 8 mm aluminum stub using silver epoxy Epo-Tek (EMS, Hatfield, PA). The mounted sample was then carefully trimmed into a smaller ∼0.5 mm^3^ tower using a Diatome diamond trimming tool (EMS, Hatfield, PA) and vacuum sputter-coated with gold palladium to help dissipate charge.

Sectioning and imaging of the sample was performed using a VolumeScope 2 SEM™ (Thermo Fisher Scientific, Waltham, MA). Imaging was performed under low vacuum conditions to introduce water vapor into the chamber, with a starting energy of 3.0 keV and beam current of 0.10 nÅ. Approximately 500 sections were cut at 50 nm thickness, providing a final 12 nm x 12 nm x 50 nm spatial resolution.

### Image Analysis and Manual Segmentation

For each fish, approximately 500 serial images encompassing an inner ear crista were denoised using Gaussian and unsharp masking filters within *Amira*. Three-dimensional reconstructions were created from manual segmentations using the brush tool in *Amira* (width=5). Five cells from three wildtype fish and five cells from three *myo7aa*^*-/-*^ mutant fish were segmented for a total of 10 cells from 6 fish. Of the total group, two wildtype and two *myo7aa*^*-/-*^ mutant cell reconstructions include the mitochondrial network and innervating neurons. Each organelle was segmented with its cellular membrane as the outer edge and was identified with organelle-specific criteria. Surface area, volume, and sphericity were calculated with the *Amira* software using the LabelAnalysis function. The following inclusion criteria for segmented cells and organelles were chosen to remove possible confounding effects of cellular damage from image preparation^16^.

#### I. Hair Cell Selection

Manual segmentation was performed only on viable hair cells which were defined as appearing healthy with no abnormal morphological features indicative of cellular death and must include stereocilia, a nucleus, mitochondria, and be innervated.

#### II. Ribbon Synapse

Segmented ribbons must present no abnormal morphology and have a clearly defined central density surrounded by tethered glutamatergic vesicles. Each ribbon synapse was segmented with its tethered glutamatergic vesicles as the outer border of the reconstruction **(Fig. S1)**.

#### III. Nucleus

Nuclei were selected during the primary search for a viable hair cell, which must show no abnormal morphology such as fragmentation or other signs of cellular death.

#### IV. Mitochondria

Mitochondria were selected for segmentation if they showed distinct cristae. The segmentation of mitochondria that showed bulging, swelling or cristolysis was done by taking the outermost boundary of the mitochondrial membrane as the edge of the segmentation. When the rupture only slightly affected the general morphology, the outer mitochondrial membrane was followed intuitively across the rupture. If the rupture caused the morphology of the mitochondrion to be completely disrupted, it was not selected for segmentation.

#### V. Afferent Nerve

The afferent nerves were identified by a large contrast in grayscale with surrounding features. Nerves were selected for segmentation if they formed a synapse with the hair cell of interest or contacted the cellular membrane. The selected nerves are not segmented in their entirety, and end approximately at the postsynaptic region.

### Innervation to Ribbon Measurement

The distance from innervation to ribbon was determined by measuring between the synaptic membrane and an ectopic ribbon’s reconstruction. This was done by first overlaying an ortho slice and ribbon reconstruction, then using the measuring tool in *Amira* **(Fig. S2)**.

### Sphericity Measurement

The ribbon reconstruction shape was quantified by sphericity, *Ψ*, which is a ratio of the surface area of a sphere with the same volume as the measured ribbon and the measured surface area of said ribbon.

Sphericity is given by the following equation^17^,

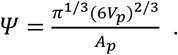

*V*_*p*_ represents the measured volume of the ribbon and *A*_*p*_ represents the measured surface area.

### Data Analysis

All data was acquired from *Amira* for EM Systems 6.7.0. Significance was determined via two-tailed two sample Student’s t-Test assuming unequal variance using Microsoft Excel.

## Results

### Three-dimensional ultrastructure of zebrafish ribbon synapses within inner ear hair cells of the cristae

To determine the effect of mutations in *myo7aa*^*-/-*^ on hair cell ultrastructure, three-dimensional reconstructions from SBFSEM images of inner ear hair cells within zebrafish cristae were created **(Fig. S1, Movie 1)**.

For both 5 dpf wildtype and mutant groups, we found that ribbon synapses are localized within the presynaptic active zone of the hair cells and adjacent to the plasma membrane **(Fig. 2)**. Consequently, the ribbons are localized around nearby regions of innervation, which are found towards the basolateral region of the hair cell. However, ribbons were also found in an ectopic position floating within the cytosol without anchorage to the basolateral membrane. Subsequently, the number and distance of ectopic ribbons to the nearest innervation were compared between groups **(Fig. S2)**. Wildtype cells contained 17 ectopic ribbons (n=5 cells; three zebrafish) with a mean (±s.e.m.) of 3.4 ± 1.4 ribbons per cell whereas the *myo7aa*^*-/-*^ mutant hair cells contained 7 ectopic ribbons (n=5 cells; three zebrafish) with a mean (±s.e.m.) of 1.4 ± 0.4 per cell (t-Test, *P* = 0.23). Wildtype ectopic ribbons had an average distance to the presynaptic plasma membrane of 455 nm ± 0.115, and the mutant ectopic ribbons had an average distance to the plasma membrane of 312 nm ± 0.116 (t-Test, *P* = 0.55). We conclude that ribbon localization occurs at the basolateral membrane near the presynaptic active region and is indistinguishable between wildtype and *myo7aa*^*-/-*^ mutant inner ear cristae hair cells.

**FIG. 2:**
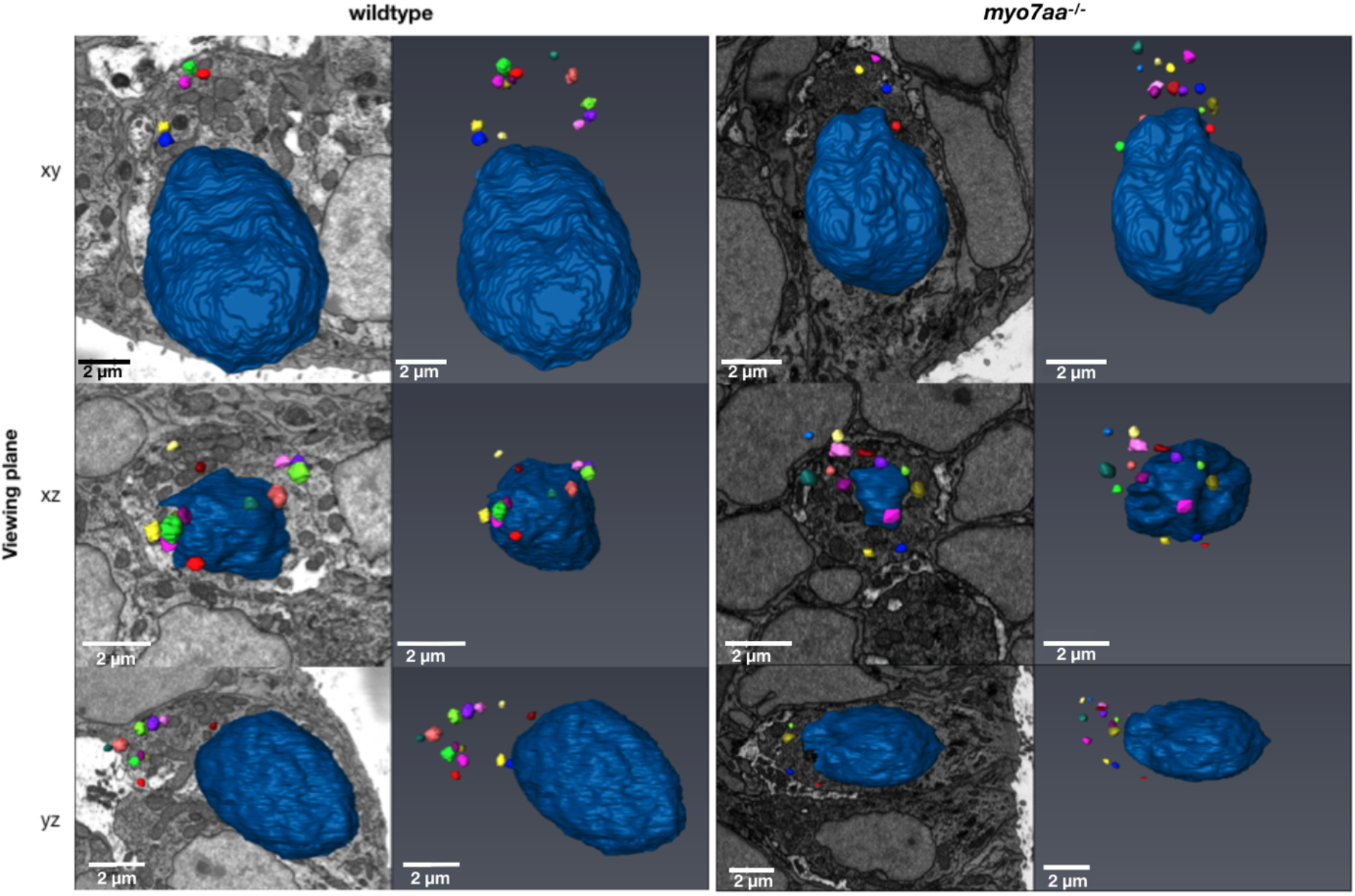
Three-dimensional reconstructions of wildtype and *myo7aa*^*-/-*^ inner ear zebrafish hair cells. Wildtype and *myo7aa*^*-/-*^ mutant hair cells are shown in three different planes (xy, xz, and yz). The ribbon synapses are individually colored and the nucleus is shown as dark blue. The leftmost column of each group shows the three-dimensional reconstruction juxtaposed with a corresponding SBFEM image. Scale Bars: 2 μm. Slight differences in length are due to small changes in zoom between planes.

Sphericity was calculated to determine differences between wildtype and *myo7aa*^*-/-*^ mutant ribbon synapse vesicle organization. No statistically significant difference was observed, with wildtype ribbon reconstructions (n=65 ribbons) having a mean sphericity of 0.855 ± 0.005 versus *myo7aa*^*-/-*^ mutant ribbon reconstructions (n=67 ribbons) having a mean sphericity of 0.847 ± 0.005 (t-Test, *P* = 0.29). These results confirmed that there is no discernible difference in shape between the wildtype and *myo7aa*^*-/-*^ mutant ribbon reconstructions **(Fig. 2, Fig. 3)**.

**FIG. 3:**
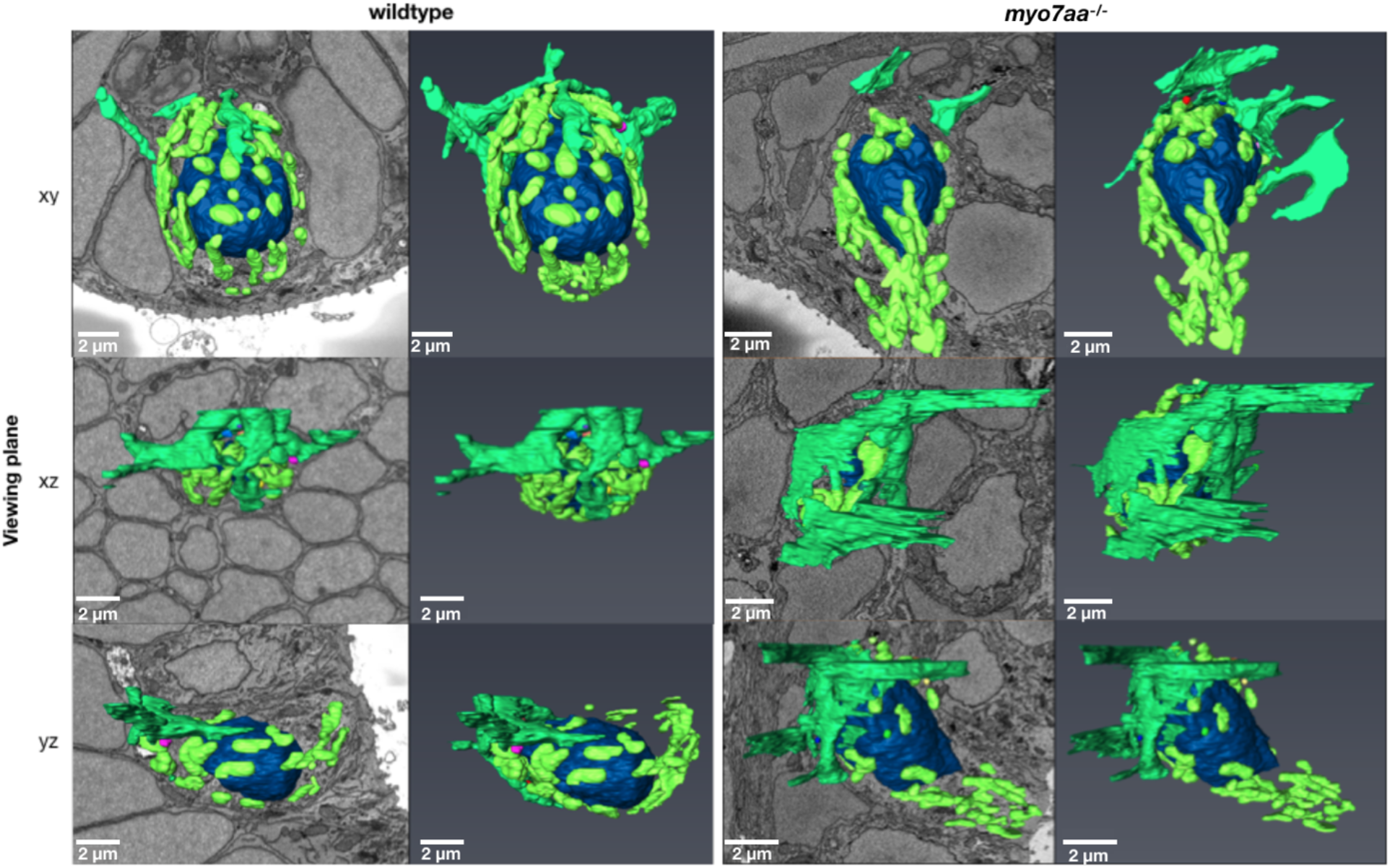
Mitochondria, nerve, ribbons, and nucleus three-dimensional reconstructions of wildtype and *myo7aa*^*-/-*^ inner ear zebrafish hair cells shown in three different planes (xy, xz, and yz). The ribbon synapses are individually colored, the nucleus is shown as dark blue, the mitochondria are neon green, and the nerve is shown as a darker green. The leftmost column of each group shows the three-dimensional reconstruction juxtaposed with a corresponding SBFEM image. Scale bars: 2 μm. Slight differences in length are due to small changes in zoom

Similarly, no significant difference in ribbon distribution was observed. Wildtype hair cells have a mean ribbon count of 13.0 ± 1.4 and *myo7aa*^*-/-*^ mutant hair cells have a mean ribbon count of 13.4 ± 2.6 (**Fig. 4D**, n=10 cells, 6 zebrafish, t-Test, *P* = 0.90).

**FIG. 4:**
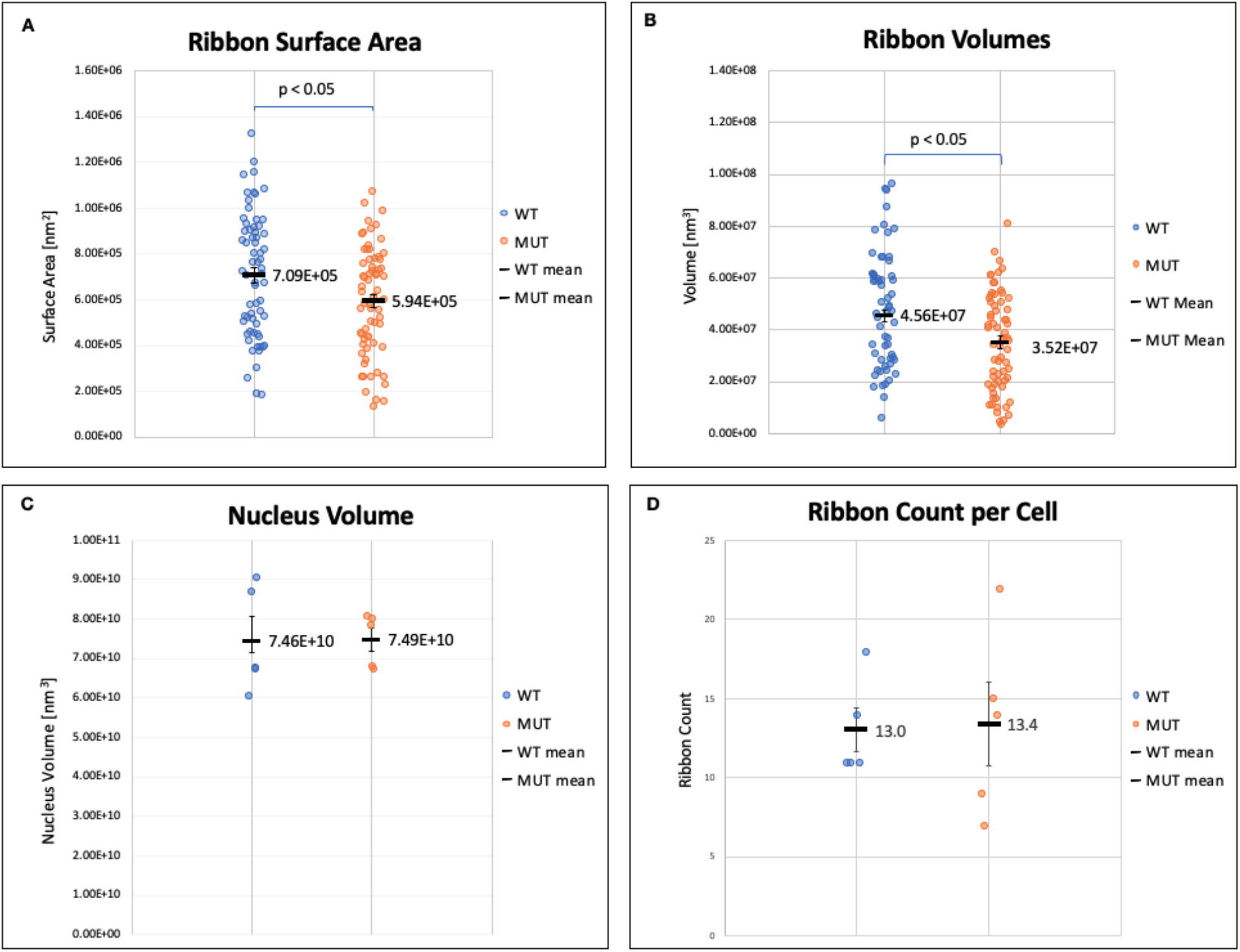
Segmentation Metrics: Dot plot representations with mean (bold black line) and standard error bars for Ribbon Surface Area **(A)**, Ribbon Volume (B), Nucleus Volume **(C)**, and Ribbon Count per Cell **(D)**.

Ribbon reconstruction volumes and surface areas were calculated to further investigate the morphology of both groups. The volume and surface area of each reconstruction includes all space within the tethered glutamatergic vesicles **(Fig. S1)**. The wildtype ribbon reconstructions (n=65 ribbons; five hair cells; three zebrafish) have an average volume of 4.56×10^7^ nm^3^ ± 2.74×10^6^ and the *myo7aa*^*-/-*^ mutant ribbon reconstructions (n=67 ribbons; five hair cells; three zebrafish) have a smaller average volume of 3.52×10^7^ nm^3^ ± 2.31×10^6^ (**Fig. 4B**, t-Test, *P* = 0.006). Similarly, wildtype ribbon reconstructions (n=65 ribbons) have an average surface area of 7.09×10^5^ nm^2^ ± 3.33×10^4^ whereas mutants (n=67 ribbons) have a smaller average surface area of 5.94×10^5^ nm^2^ ± 2.88×10^4^ (**Fig. 4A**, t-Test, *P* = 0.01). Thus, *myo7aa*^*-/-*^ mutant ribbon reconstructions have a statistically lower volume and surface area compared to wildtype.

### Inner ear hair cell mitochondria three-dimensional morphology

Mitochondria were segmented to qualitatively compare three-dimensional structure between wildtype and *myo7aa*^*-/-*^ mutant hair cells **(Movie 2, Movie 3)**. In both wildtype and mutant cells, the mitochondria form a complex tubular network extending from the synaptic region of the cell to stereociliary sites with a lower mitochondrial density between these two regions. Also, the mitochondria show a diversity of morphological forms **(Fig. 3)**.

Although uncommon, mitochondrial swelling, bulging and cristolysis were found in both wildtype and mutant hair cells. Often, the outer membrane of the abnormal mitochondria remained intact. However, some mitochondria, in both wildtype and mutants, showed ruptured outer membranes **(Fig. 5)**. To determine if a true phenotype exists, further study is required on a larger number of hair cells.

**Fig. 5:**
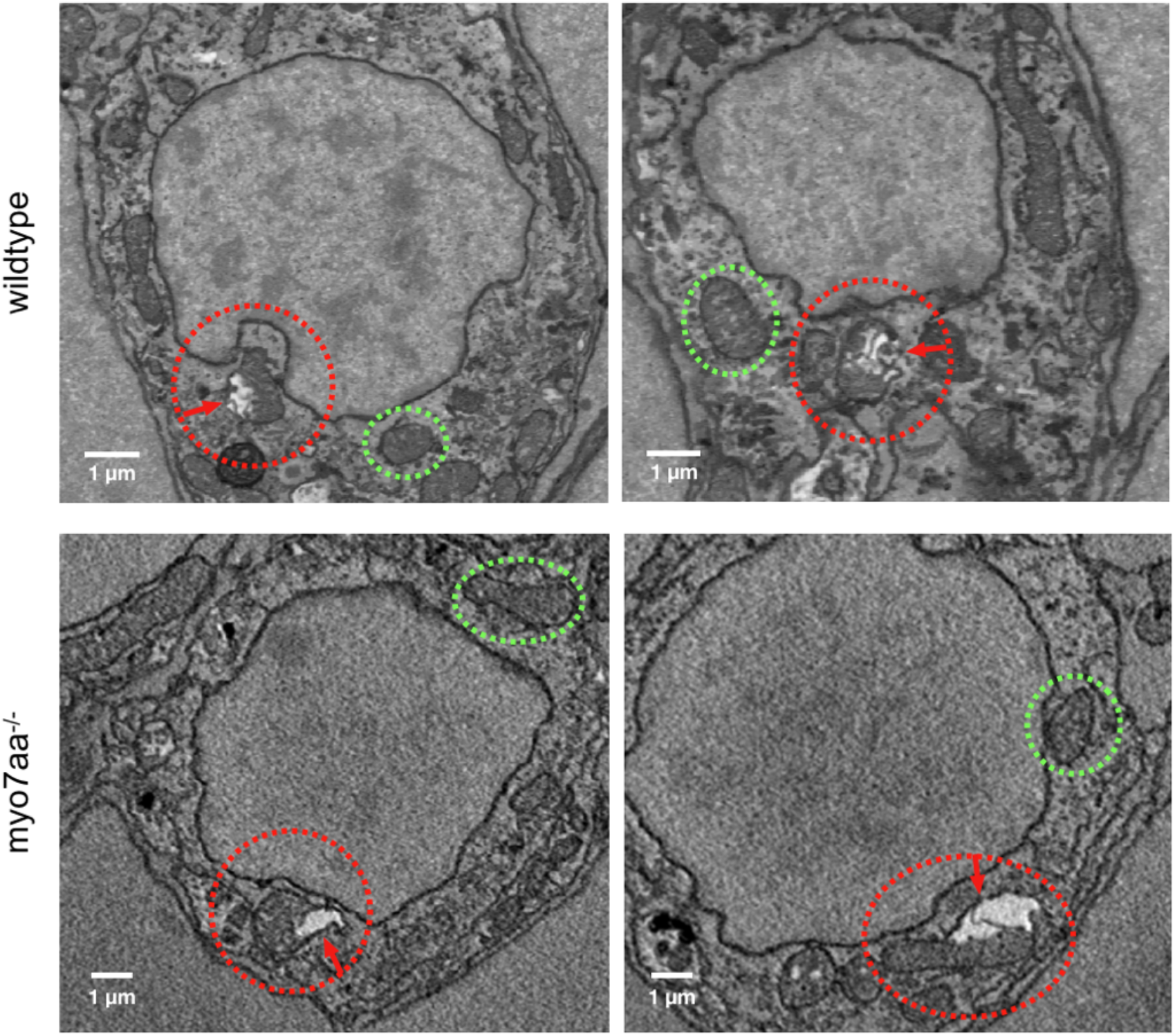
Mitochondrial Abnormalities: SBFSEM images of wildtype and myo7aa-/- inner ear hair cells. Normal mitochondria are circled in green. Mitochondria with abnormal characteristics such as bulging, cristae disruption, and swelling were found in both groups and are circled in red with an arrow indicating the region which shows abnormal properties.

### Qualitative analysis of nerve morphology

Observationally, both the wildtype and mutant inner ear hair cells are innervated and have basolateral synapses which are complex and far-reaching. Nerve fibers form extensive calyx-like postsynaptic connections, which cover large surface areas surrounding nearly half of the hair cell in which they innervate **(Fig. 3)**. In contrast to simple boutons, these synaptic connections are highly branched and form small projections. Furthermore, within the *myo7aa*^*-/-*^ mutant hair cells, the synapse takes on a more planar form. Finally, in both models, it can be seen that each inner ear hair cell may be innervated by more than one neuron extending from the basal layer.

## Discussion

This study was conducted to uncover the three-dimensional morphology of the ribbon synapses, mitochondrial networks, and synaptic neurons within inner ear crista of wildtype and *myo7aa*^*-/-*^ mutant zebrafish which, to date, is not well characterized. This was accomplished via SBFSEM imaging and analysis with the *Amira* software package.

We found no difference in ribbon reconstruction localization and shape between wildtype and *myo7aa*^*-/-*^ mutants. This agrees with previous findings that suggest ribbon localization, development and stabilization is dependent upon the presence of innervation rather than the lack of mechanotransduction or synaptic activity^14^. Furthermore, previous studies show ribbon size and morphology are regulated during development, 1-4 dpf, by spontaneous action potentials mediated by L-type calcium channel activity and regulated by mitochondrial calcium uptake^18-21^. It has been shown that mature mutant zebrafish which lack mechanotransduction display longer intervals of spontaneous calcium activity and longer periods of inactivity^19^.

However, at maturation, the effects of calcium influx on ribbon size and morphology are remarkably reduced^18,21^. Hence, our study demonstrates no difference between, 5 dpf *myo7aa*^*-/-*^ mutants and wildtype hair cells for presynaptic localization or ribbon vesicle organization, indicating that synaptogenesis may be unabated by mutations in *myo7aa*^*-/-*^.

Ribbon counts were found to be comparable. Interestingly, we report wildtype hair cells with an average of 12.0 ribbons per hair cell and mutants with an average of 12.4 ribbons per hair cell. These large inner ear ribbon counts strikingly contrast those of neuromast hair cells, which only average 3-5 single ribbons^22^. Moreover, it has been previously shown that neuromast hair cells in *myo7aa*^*-/-*^ mutants have on average two ribbons per hair cell whereas wildtype cells have three^13^. We may expect this same effect to be evident within the inner ear cristae as well, however, our results show a slight increase in ribbon counts within *myo7aa*^*-/-*^ mutant cells. However, conclusive interpretation of this result will require a larger sample size.

The number of ectopic ribbons and their distance to the synaptic membrane were comparable between groups. This may further indicate that ribbon development is unaffected by the lack of mechanotransduction within *myo7aa*^*-/-*^ mutants, as ectopic ribbons may be a sign of altered development or irregular turnover^14,23^. However, we found the wildtype ribbon reconstructions to have a statistically higher volume and surface area compared to the *myo7aa*^*-/-*^ mutant reconstructions. The three-dimensional reconstructions presented are of the tethered glutamatergic vesicles of the ribbon synapse **(Fig. S1B)**. As a result, multiple factors may contribute to the disparity between volume and surface area measurements of wildtype and mutant reconstructions. Previous work has demonstrated differences in vesicle tethering between wildtype and *ribeye b-EGFP* transgenic ribbons, with wildtype ribbons presenting a longer tethering distance^24^. Within our study, a similar tethering effect may exist, an effect of ribeye protein morphology, or a change in tethered vesicle density. Elucidating the cause for change in ribbon reconstruction volume will require further investigation.

Additionally, mitochondria and innervating neurons were reconstructed for qualitative analysis. The mitochondrial network showed a clear localization near the presynaptic active region of the cell, and near the stereocilia as previously reported in neuromasts^17^. This localization pattern could be reasonably attributed to the high energy demands of calcium-dependent stereocilia mechanotransduction and of neurotransmission at the presynaptic active region^24^. Abnormal mitochondria, which presented swelling and cristolysis, were found in both wildtype and *myo7aa*^*-/-*^ mutants. Observationally, these features seem to be indicative of apoptosis, mitophagy, oncogenic stress or hypoxia rather than mitochondrial fusion or fission^26-29^. Causality of these abnormal mitochondria is an area for future studies.

As reported previously, the innervating neurons show highly branched calyx-like synapses in both wildtype and *myo7aa*^*-/-*^ mutant groups^22^. Prior work in lateral line neuromasts showed that each sensory hair cell associates with multiple afferent terminals, and ribbons form connections with two or more of these terminals creating a complex microcircuitry^7^. It is likely the redundant circuitry seen in the neuromasts is also seen within the inner ear and is responsible for the complex innervation we find within the crista. However, further segmentation extending past the basal layer of the crista in all neurons would be required to investigate this. We also find that the mutant neuron reconstruction appears more planar than the wildtype, covering a larger surface area of the hair cell. This may be a compensation effect, as the *myo7aa*^*-/-*^ mutants fail to depolarize in response to stimuli.

## Limitations

The SBFSEM imaging and manual segmentation process is labor intensive^7^. Also, image quality and greyscale may vary between samples. These constraints forced our study to a small number of cells, reducing the statistical power.

## Conclusion

In all, our study shows that the utilization of SBFSEM and three-dimensional reconstruction is a promising method to support experimental results and will advance the field by providing critical new tools for hypothesis generation.

## Author Contributions

Conceptualization (KCR, AK, LAS, JD); Data Curation (KCR, TAC, JD, LAS); Formal Analysis, Investigation, Validation, Writing – review and editing (KCR, AK, JD, SAC, TAC, LAS); Funding Acquisition (LAS); Methodology (KCR, JD, TAC); Project Administration (KCR, LAS); Resources (TAC, LAS); Software (KCR, AK, JD, TAC); Supervision (LAS); Visualization (KCR, JD, TAC); Writing - original draft (KCR).

## Data Availability

Contact Kenneth C. Riley Jr., and Lisa A. Schimmenti for raw data files.

