## Supplemental Figures for "Three-dimensional structure of inner ear hair cell mitochondrial networks and ribbon synapses in a zebrafish model of Usher syndrome type 1B"

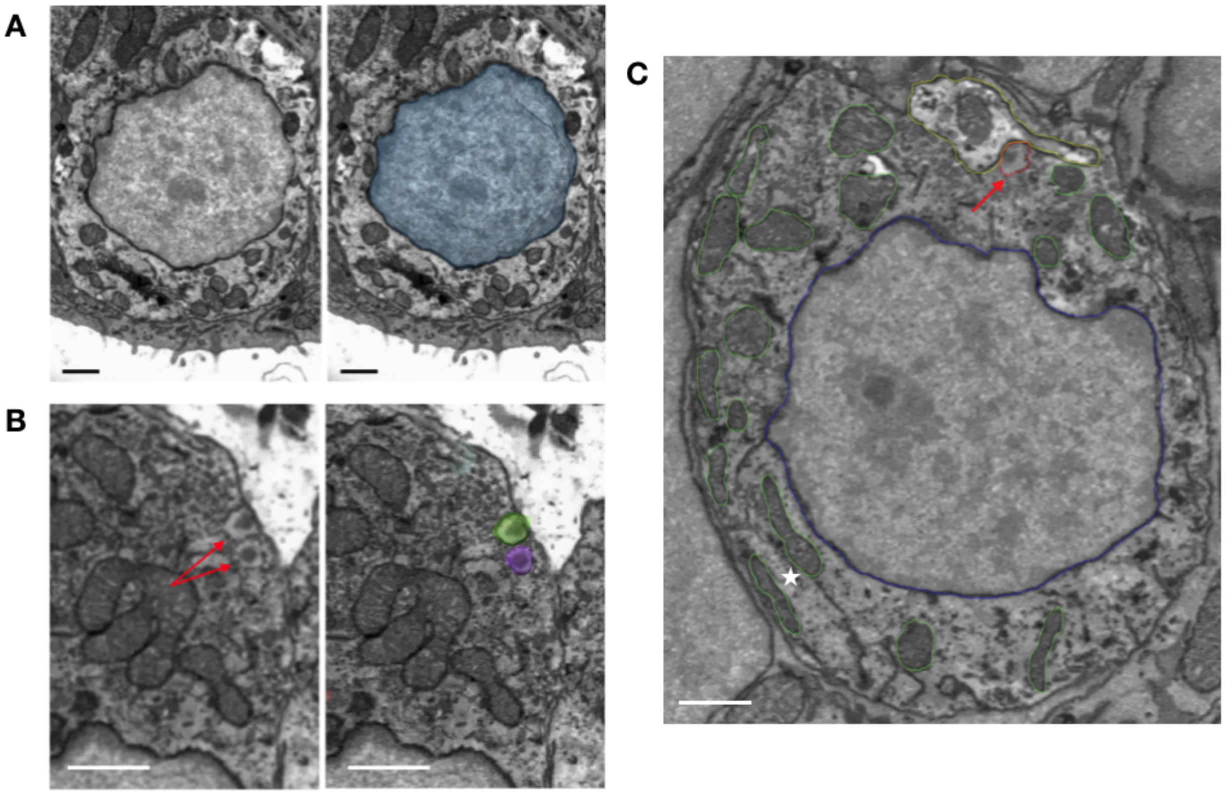


**S1. *Amira* Image Segmentation and Measurement:** A) Single wildtype SBFSEM hair cell section image before (left) and after nuclear segmentation. The nuclear membrane is traced to segment the structure. Hundreds of section images are individually segmented to generate three-dimensional reconstructions. Scale Bars: 1µm. B) Identical SBFSEM images to demonstrate ribbon synapse segmentation. Ribbon synapses prior to segmentation are indicated with red arrows in the left panel. The right panel, after segmentation, shows that the ribbon reconstruction follows the tethered glutamatergic vesicles. Scale Bars: 1 µm. C) SBFSEM section image of a single wildtype inner ear hair cell with full segmentation, indicated by colored outlines, including the nucleus (blue), the mitochondria (green), a ribbon synapse (red), and an innervating neuron (yellow). The red arrow highlights a segmented ribbon, and the star highlights an example of segmented mitochondria. Scale Bar: 1µm.

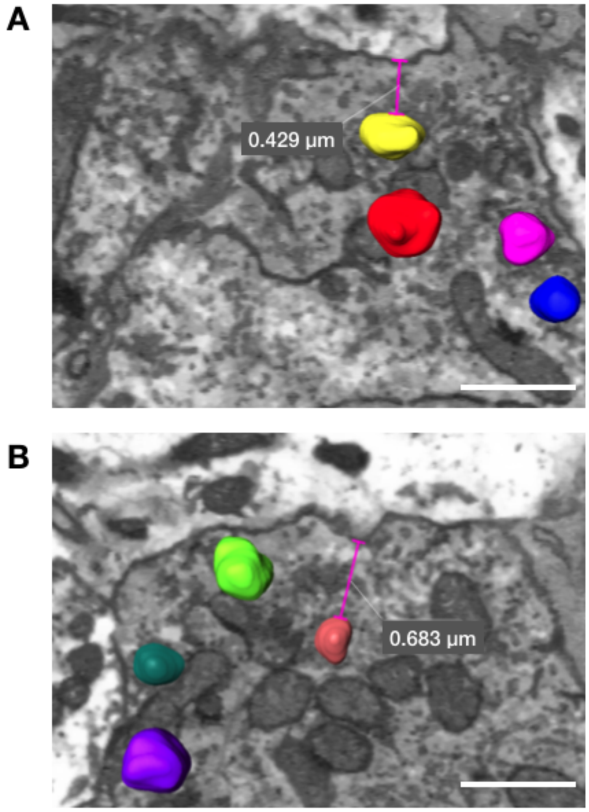


**S2. Innervation to Ribbon Distance:** Measurement between reconstructed wildtype ectopic ribbons and the presynaptic membrane in two individual SBFSEM section images in *Amira* (A, B). Scale Bars: 1µm.

### Movie Legends

**Movie 1:** Wildtype nucleus and ribbon three-dimensional reconstruction with corresponding SBFSEM images. The nucleus is slightly transparent and shown in red. Ribbons are individually colored. Created with *Amira* animation tools**.**

**Movie 2:** Wildtype hair cell reconstruction including the nucleus (blue), ribbon synapses (individually colored), mitochondria (neon green), and nerve (dark green). Created with *Amira* animation tools**.**

**Movie 3:** *myo7aa*$^{-/-}$ mutant hair cell reconstruction including the nucleus (blue), ribbon synapses (individually colored), mitochondria (neon green), and nerve (dark green) with corresponding SBFSEM images. Created with *Amira* animation tools**.**
